# Magnetic Resonance Imaging Tracking of Distal Inflammatory Changes following Mild Traumatic Brain Injury (mTBI) in rats

**DOI:** 10.1101/057679

**Authors:** SB Gillham, JD Figueroa, B Bartnik

## Abstract

Sensorimotor disturbances continue to represent one of the most debilitating and widely reported complications in patients suffering mild traumatic brain injuries (mTBI). Loss of peripheral neuronal function at sites distal and disconnected to the central nervous control central centers is well documented. Distal muscular atrophy, complex regional pain symptoms, and poor wound healing are just a few of the many complications with often more severe secondary complications such decubitus ulcers and osteomyelitis seen at sites in the body distal to the center of injury. MRI has been widely established as a diagnostic and therapeutic planning tool in patients and animal models with neuronal disease. However, studies investigating the neural correlates of spinal cord changes after TBI are lacking. Here, we used T2 MR imaging to determine the effects of mTBI on the morphology and inflammatory changes of the spinal cord. We hypothesize that rats receiving mTBI utilizing a controlled cortical impact (CCI) contusion will demonstrate T2 signal changes at distal locomotor centers in the spine. Experimental mTBI and sham groups of Sprague-Dawley rats were used (n = 2 sham and 4 experimental). A mild CCI was applied to the right brain cortex. Rats were sacrificed at 60 days post injury and spinal cords harvested for ex vivo MRI T2 analysis. Focal areas/lesions of increased T2 hyperintensity were noted in mTBI injured rats (n = 4).

Experimental group of rats also demonstrated secondary spinal cord locomotor and sensation adverse effects clinically. MR imaging showed volumetric reductions and T2 signal changes in the cervical, thoracic, and lumbar segments of the spinal cord at 8 weeks’ post-injury. T2 intensity values were elevated in all experimental groups in comparison to the sham group within the distal cord, suggesting that remote CCI causes secondary spinal cord inflammation and neurodegeneration at distant sites. These findings also further support the idea that the most peripheral nerves and spinal cord will be most negatively affected by a TBI. While our research is in its preliminary stages, our results further confirm that mTBI has more far reaching effects than previously understood. T2 MRI is an effective tool to assess the extent of spinal cord injury related to antecedent TBI.

## Introduction

Mild traumatic brain injury (TBI) is well known to lead to substantial socioeconomic cost and serious secondary consequences^1–3^. Although mTBI is often associated with good prognosis, it remains one of the leading causes of disability and death among children and young adults. TBI represents a huge economic burden to our society, with estimated total annual cost of 76.5 billion ^4,5^. Sensory and motor disturbances are among the most debilitating consequences of mTBI, resulting in longer hospital stays and delayed recovery during rehabilitation.

It is unclear how a cortical concussive injury can affect distant spinal cord centers. Studies investigating how inflammation and maladaptive plasticity compromise spinal cord integrity at sites distant to the initial TBI are almost nonexistent. Zing et al., recently reported that TBI reduced levels of critical “molecular systems important for synaptic plasticity (BDNF, TrkB, and CREB) and plasma membrane homeostasis (4-HNE, iPLA2, syntaxin-3) in the lumbar spinal cord” ^6^. Chen et al., showed that there are more distant neurodegenerative changes at sites uninvolved by the initial TBI, including increased neuronal degeneration and inflammation involving the ipsilateral hippocampus and thalamus between sham and injured groups who had received a mild controlled cortical contusion (CCI)^7^. It was also reported that localized BBB extravasation causes local and more distant local degeneration and likely induces a longer lasting inflammatory response ^7^. Currently, there are no efficient treatments to treat the detrimental functional deficits associated with TBI.

Recent large review studies are now accepting the role of omega-3 polyunsaturated fatty acids (ω-3 PUFAs) in conferring multiple health benefits and decrease the risk of neurological disorders^8,9^. PUFA’s are vital for neurodevelopment and regeneration. Epidemiological studies have suggested that neurotrauma reduces the levels of docosahexaenoic acid (DHA), a potent anti-inflammatory dietary-essential fatty acid also implicated in neural function. Furthermore, previous research has suggested that, Docosahexaenoic acid (DHA, C22:6 n-3) is neuroprotective when administered following and before SCI and TBI^9,10^. DHA pretreatment significantly increased the percentage of white matter sparing and resulted in axonal preservation.

Prior animal studies have shown that pre-treated animals exhibited lower sensory deficits, autonomic bladder recovery, and early improvements in locomotion that persisted for at least 8 weeks after spinal cord trauma following administration of PUFA’s. ^5,10,11^

Here, we investigated the potential neural basis for the disruptive sensorimotor effects of brain injury following mTBI in rats. We hypothesize that rats receiving mTBI utilizing a controlled cortical impact (CCI) contusion will demonstrate T2 signal changes at distal locomotor centers in the spine. The current study employs T2 MR imaging to determine the impact of mTBI on the spinal cord morphology. Once T2 MR imaging has been validated as a viable tool to assess inflammatory changes at remote sites to the initial injury, we can then use MR imaging as a tool to assess the neuroprotective effects of PUFA’s on the more distant central and peripheral nervous system.

## Methods

Animals: Experimental mTBI and sham groups of Sprague-Dawley rats were used (n = 2 sham and 4 experimental).

TBI model (CCI): Animals were deeply anesthetized with a mixture of ketamine (80⁏mg/kg) and xylazine (10⁏mg/kg). A parietal craniotomy was performed and a mild controlled cortical injury was then administered (CCI; 4⁏mm diameter tip, 0.5⁏mm depth, 6.0⁏m/s speed, 200⁏ms dwell). The dura was left intact. The sham animals received only a parietal craniotomy to expose the dura.

During surgery, the body temperature of the animals was maintained at 37°C using a thermostatically controlled heating pad with a rectal thermometer (Physitemp TCAT-2LV; Physitemp, Clifton, NJ). After surgery, the epidermis and subcutaneous tissues were closed with wound clips. Immediately after injury, the skin incision was closed with nylon sutures, and 2% lidocaine jelly was applied to the lesion site to minimize any possible discomfort.

Animals were allowed to survive for 8⁏weeks post operatively. Rats were then sacrificed and spinal cords harvested for ex-vivo MRI T2 analyses at 60 days post injury or sham. Spinal cords were suspended in a highly crossed linked Carbopol^®^ 974P NF Polymer which has been previously shown to yield superior signal to noise ratios ^12^.

MR Imaging: Utilizing an 11.7T animal MRI scanner, T2 intensity values (2d sequence; TR 3594 ms; TE 10.2 ms; NEX 4; FOV 1.5 cm; Matrix 256; 30 slices; 1 mm slice thickness; 0.5 mm slice interval) were measured at 9 different locations throughout each spinal cord to measure signal change within the dorsal column white matter (cervical level/above the lesion, mid thoracic level at the level of the lesion, and distal spinal cord below lesion in the region of the upper lumbar spine). T2 values were measured at level of cord lesions in the experimental group and at the equivalent level in the sham group of rats which showed no evidence of a focal T2 hyperintense lesion (*Figure 1*). A single T2 hyperintense mid thoracic lesion was identified all experimental groups of rats. DTI imaging was also performed for further research.

**Figure 1:**
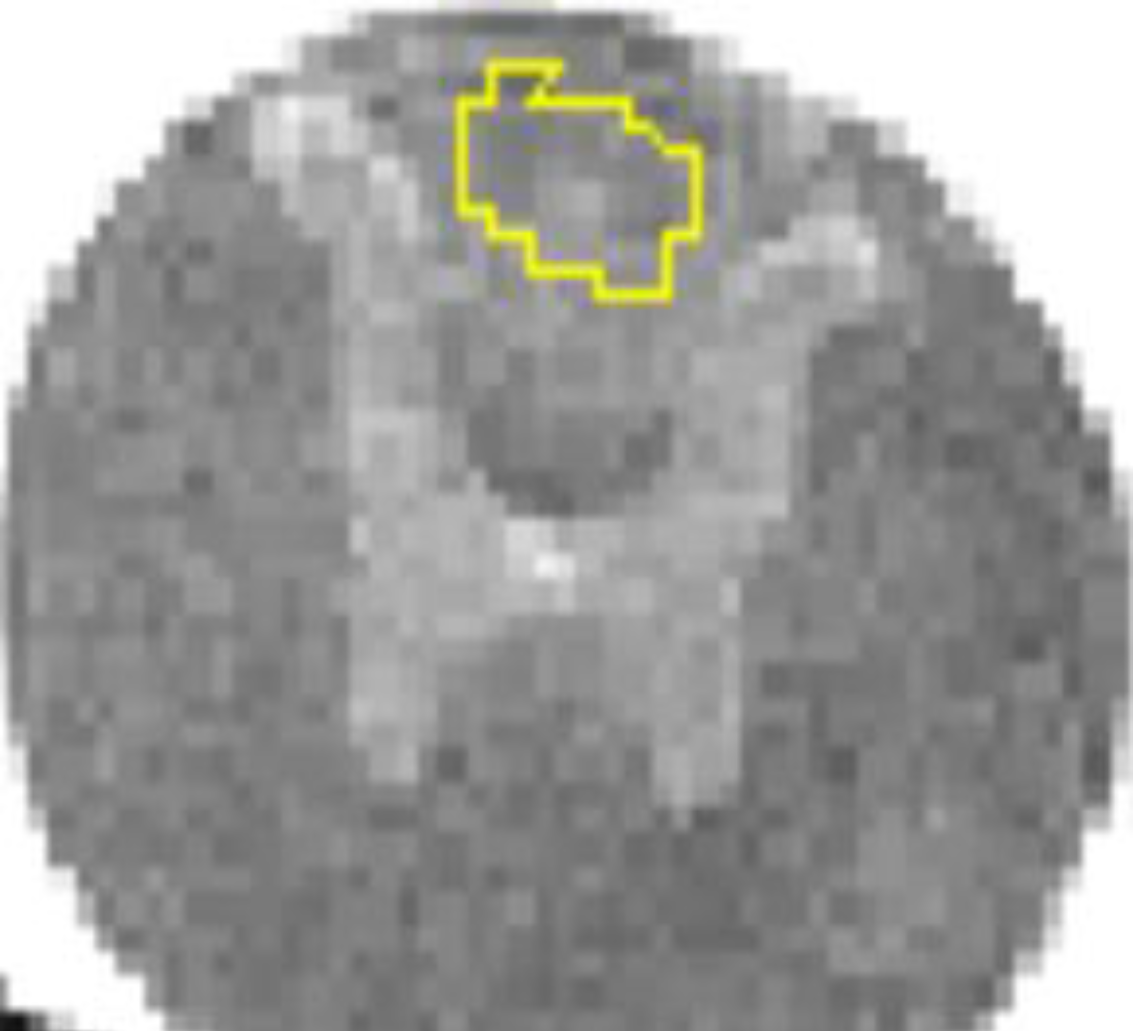
Example of T2 signal intensity measurement obtained in the thoracic spine in the expected location of the dorsal columns.

Statistical Methods: To measure the distribution of T2 intensity values in the dorsal column white matter, statistical analysis was performed using normalized Kolmogorov-Smirnova. Group differences were measured using de-trended Shapiro-Wilk tests.

## Results

Focal areas of increased T2 signal (T2 hyperintense lesions) were noted in all spinal cords of the experimental group (n=4) (*Figure 2*). This same group of experimental rats also demonstrated secondary spinal cord locomotor and sensation adverse effects clinically which will be published in further research and are not discussed within the context of this current paper.

**Figure 2:**
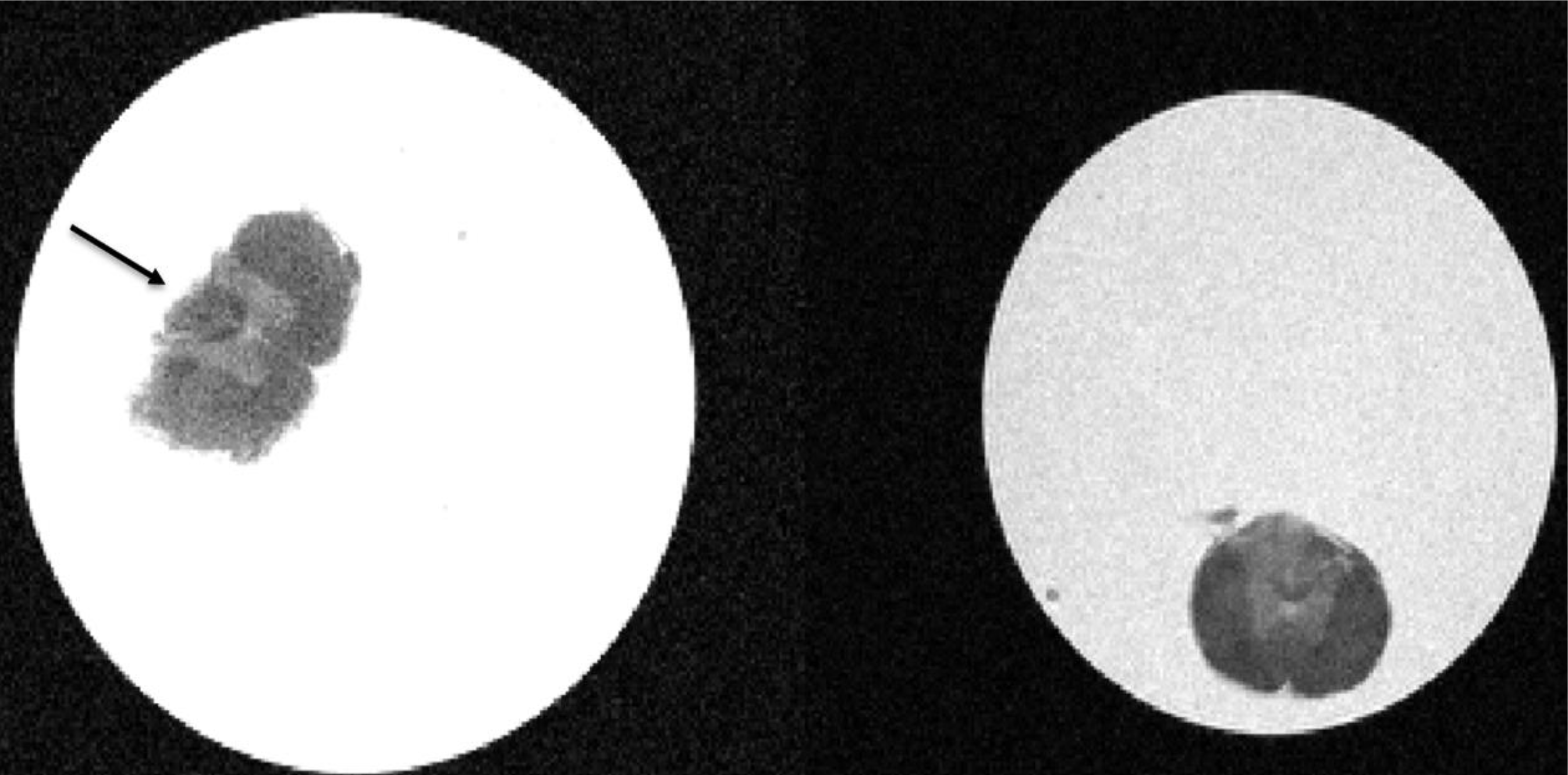
T2 hyperintense lesions were noted in 4 out of the 6 cords at the level of the mid thoracic spine.

T2 hyperintensity values in the cervical cord and level of the lesions demonstrated normal random T2 value distribution on both normal and de-trended Q-Q plots of both sham and experimental groups (*Figure 3*). T2 intensity values were elevated and showed loss of the normal random T2 value distribution (*Figure 4*) in the experimental groups in comparison to the sham group within the distal cord below the level of the lesion (*Figure 5*).

**Figure 3:**
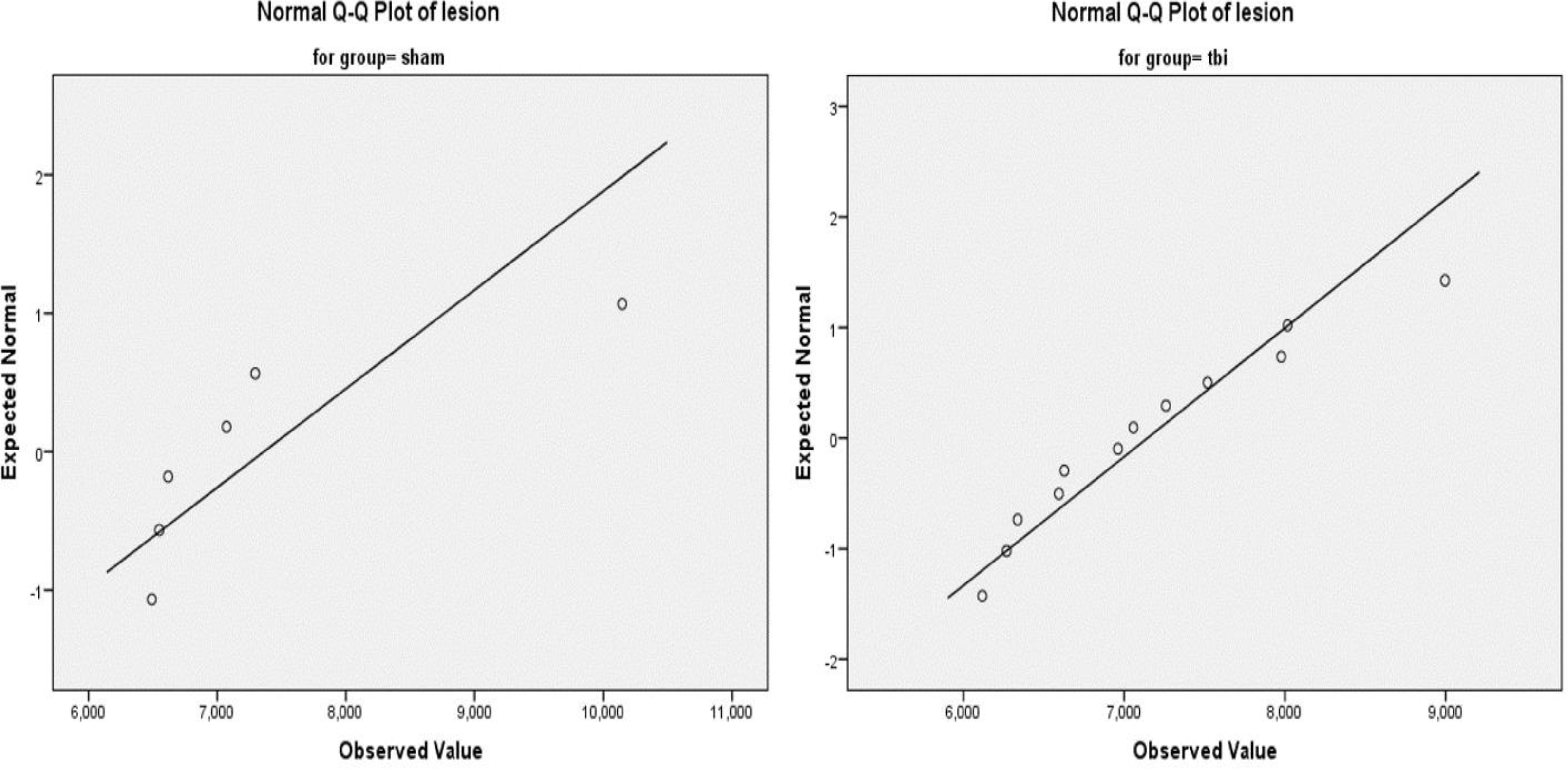
T2 values of sham and mTBI groups in the cervical (above lesion) and thoracic cord (at lesion) demonstrate expected normal random distribution.

**Figure 4:**
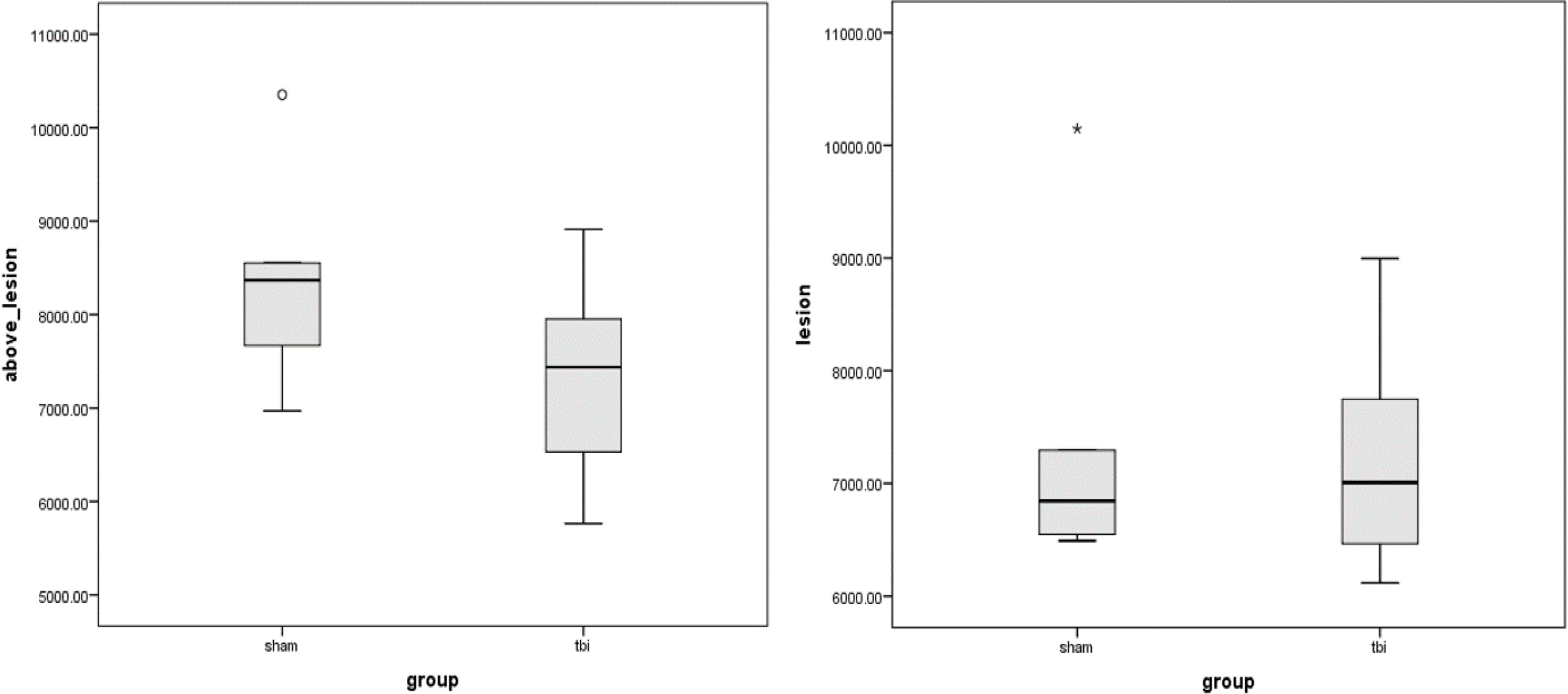
Increased distribution of quantitative T2 values in the mTBI group was identified throughout the spine.

**Figure 5:**
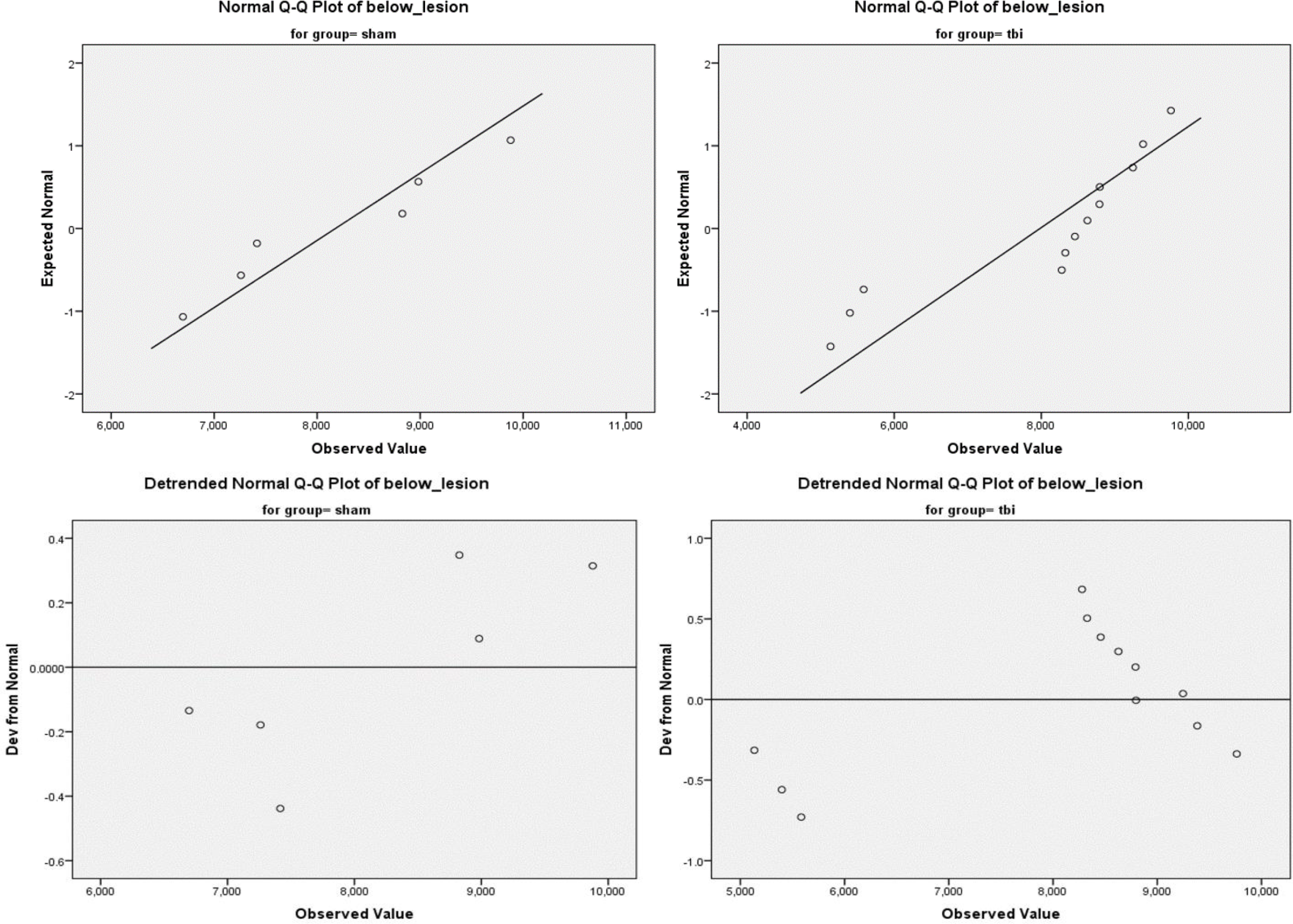
T2 values of sham and mTBI groups in the distal cord below the level of the lesion demonstrates significantly increased T2 values.

## Discussion

Our findings indicate remote CCI causes secondary spinal cord inflammation and neurodegeneration at distant sites. Our findings also reinforce that the most peripheral nerves and spinal cord will be most negatively affected by a TBI and that MRI is a viable tool to assess extent of distant neurodegeneration and may be used in the future as a predictive tool.

While prior studies have directly shown decreased volumes and increased inflammation in the ipsilateral slightly more local regional structures such as the hippocampus and thalami^7^, there has been little researching showing distant spinal cord inflammation and/or degeneration to account for the frequently observed secondary medical complications seen in many TBI patients. Our research remains in its preliminary stages; however, our results further support that mTBI has more far reaching effects than previously understood. To our knowledge, this is the only research that has demonstrated remote lesions in the spine following mTBI. This may further help explain the impaired synaptic plasticity and reduced plasma membrane homeostasis in the lumbar spine as recently reported^6^. This has significant implications for future research and understand the distal muscular atrophy and complex regional pain symptoms often seen with patients following TBI. In addition, identifying focal T2 hyperintense lesions within the spine could have significant impact on targeted treatment strategies for patients with distal locomotor symptoms. T2 MRI has been long used to identify direct spinal cord injury, but has not been used as a tool for assessing remote injury related to TBI. Our research supports T2 MRI as an effective tool in assessing the extent of spinal cord injury related to antecedent TBI, which has not been previously established. This could have profound implications in categorizing and providing some measure of staging for severity and expected outcomes of patients with TBI.

With this additional supporting evidence that MRI T2 imaging can accurately assess distal spinal cord injury, our group plans to conduct further research demonstrating MRI’s ability to document improved cord inflammation in experimental groups receiving omega-3 PUFA enriched rat chow following TBI. The secondary changes to the spinal cord related to TBI are thought to be responsible for at least some of the secondary consequences of TBI. We will assess for T2 signal changes in the cervical, thoracic, and more distal segments of the spinal cord at 8 weeks post-injury ^13^.

We hypothesize that diets enriched with omega-3 PUFAs will ameliorate the observed signal changes in the spinal cord. This study may have implications for the management and rehabilitation of patients after mTBI, suggesting a strategy to minimize distal spinal cord secondary consequences through the use of omega-3 PUFAs.

